# Occipital Intralobar fasciculi and a novel description of three forgotten tracts

**DOI:** 10.1101/2020.07.07.191767

**Authors:** Maeva Bugain, Yana Dimech, Natalia Torzhenskaya, Michel Thiebaut de Schotten, Svenja Caspers, Richard Muscat, Claude J Bajada

**Affiliations:** Department of Physiology and Biochemistry, Faculty of Medicine and Surgery, The University of Malta; Department of Cognitive Sciences, Faculty of Media and Knowledge Sciences, The University of Malta; Brain Connectivity and Behaviour Laboratory, Sorbonne Universities, Paris, France; Groupe d’Imagerie Neurofonctionnelle, Institut des Maladies Neurodégénératives - UMR 5293, CNRS, CEA University of Bordeaux, Bordeaux, France; Institute of Neuroscience and Medicine (INM-1), Research Centre Juelich, 52425, Juelich, Germany; Institute for Anatomy I, Medical Faculty, Heinrich-Heine-University Duesseldorf, 40221, Duesseldorf, Germany

**Keywords:** White Matter, Dejerine, Anatomy, Tractography

## Abstract

The continuously developing field of magnetic resonance imaging (MRI) has made a considerable contribution to the knowledge of brain architecture. It has given shape to a desire to construct a complete map of functional and structural connections. In particular, diffusion MRI paired with tractography has facilitated a non-invasive exploration of structural brain anatomy, which has helped build evidence for the existence of many association, projection and commissural fiber tracts. However, there is still a scarcity in research studies related to intralobar association fibers. The Dejerines’ (two of the most notable neurologists of 19^th^ century France) gave an in-depth description of the intralobar fibers of the occipital lobe. Unfortunately, their exquisite work has since been sparsely referred to in the modern literature. This work gives the first modern description of many of the occipital intralobar lobe fibers described by the Dejerines. We perform a virtual dissection and reconstruct the tracts using diffusion MRI tractography. The virtual dissection is guided by the Dejerines’ treatise, *Anatomie des Centres Nerveux.* As an accompaniment to this article, the authors provided a French-to-English translation of the treatise portion concerning intra-occipital fiber bundles. This text provides the original description of five intralober occipital tracts, namely: the stratum calcarinum, the stratum proprium cunei, the vertical occipital fasciculus of Wernicke, the transverse fasciculus of the cuneus (TFC) and the transverse fasciculus of the lingual lobe of Vialet. It was possible to reconstruct all these tracts except for the TFC – possibly because its trajectory intermingles with a large amount of other fiber bundles. For completeness, the recently described sledge runner fasciculus, although not one of the Dejerine tracts, was identified and successfully reconstructed.

## INTRODUCTION

The advent of novel *in vivo* magnetic resonance imaging (MRI) techniques has heightened the desire to construct a comprehensive atlas of the human brain connective network – the so-called human connectome (Van Essen and Ugurbil 2012; Shibata et al 2015; Sotiropoulos and Zalesky 2019). In humans, diffusion MRI paired with tractography permits a non-invasive three-dimensional reconstruction of large-scale white matter system organisation, which can then be compiled into an atlas (Wakana et al 2004) (see review: (Jbabdi and Behrens 2013; Jbabdi et al 2015; Jeurissen et al. 2017). Diffusion MRI has contributed to the verification of many association fiber pathways (Meola et al 2015; Bao et al 2017), as well as to the redefinition of their boundaries and relations (Kamali et al 2014; Sampson et al 2015; Hau et al 2016; Wu et al 2016; Weiner et al 2017; Panesar et al 2019; Schurr et al 2019). Recent success in discovering novel or forgotten association fibers has stoked interest in the less conspicuous short-range fiber connections, which can be more challenging to delineate (Yeatman et al 2014; Uesaki et al 2018; Koutsarnakis et al 2019; David et al 2019).

Despite the rapid advancement in the technology and the connectome mapping techniques available, there is a paucity in research related to short-range association fibers in the occipital lobe. The initial focus of this new field was instead geared towards the more prominent, and seemingly more physiologically crucial, long-range association fibers. An additional issue with MRI-based tractography studies is that the findings tend to remain speculative, devoid of tangible experimental evidence and ground truth, unlike those of axonal tracer studies (Donahue et al 2016).

To partially overcome the limitations inherent in diffusion tractography, contemporary researchers can guide their findings through post-mortem dissection studies, by consulting the tract tracer literature, or by consulting historical neuroanatomy texts. The 19^th^-century neurologists were masterfully skilled in gross dissection and fastidious in the documentation of their composite observations. As such, their findings remain relevant and still permeate today’s literature. Numerous diffusion tractography studies have successfully used classical dissections to interpret imaging results and identify false positives or false negatives, and have been successful in building on the groundwork made by their predecessors (Schmahmann and Pandya 2006; Catani et al 2012; Forkel et al 2014; Vergani et al 2014; Yeatman et al 2014). Pioneering texts have also often been an initial source of inspiration for explorative neuroanatomical studies (Forkel et al 2014; Yeatman et al 2014). Indeed, the authors had previously studied dissections by 19^th^-century neurologists Joseph and Augusta Dejerine with a view to enriching the field’s knowledge and perspectives on long-range association fiber anatomy (Bajada et al 2017). It showcased the pertinence of these historical studies and prompted further examination into the Dejerines’ other work on association short-range occipital fibers.

The 1900s was a century particularly rich in pivotal studies and debates between different schools of thoughts. One of the biggest divisions across the neuroanatomical field towards the end of the 19^th^ century was the definition and categorisation of association fibers. Meynert was one of the most prominent authorities on subcortical anatomy at the time. He postulated that true association fibers only projected in an anterior-posterior direction (Meynert 1892; Yeatman et al 2014). This was contested by Wernicke, who argued for the existence of dorsal-ventral projecting association fibers, particularly in the occipital region (Wernicke 1876; Yeatman et al 2014). Other neurologists, like the Dejerines and Sachs, supported the school of thought of Wernicke by acknowledging the existence of the vertical occipital fasciculus (VOF) (Sachs and Wernicke 1892; Déjerine and Déjerine-Klumpke 1895). Joseph Jules Dejerine (1849–1917) and his wife, Augusta Dejerine-Klumpke (1859–1927) frequently collaborated and are both individually celebrated as two of France’s most renowned 19^th^-century neurologists (Bogousslavsky 2005; Shoja and Tubbs 2007; Bajada et al 2017). Together, they pioneered anatomo-functional studies of the cerebral association fiber tracts and challenged the period’s paradigm of brain organisation (Shoja and Tubbs 2007; Krestel et al 2013; Bajada et al 2017). Arguably, the Dejerines’ seminal work is their two-volume neuroanatomical treatise, *Anatomie des Centres Nerveux*, originally published in 1895 and 1902 and then reprinted in 1980 by Mason-Elsevier (Déjerine and Déjerine-Klumpke 1895; Dejerine and Dejerine-Klumpke 1902). The Dejerines’ impressive histological sections, accompanied by beautifully intricate illustrations by H. Gillet, are relevant even in this technological era, and are thus referred to for comparison to new anatomical findings (Chou et al 2009; Catani et al 2012). Of particular interest to the authors, the Dejerines dedicated a portion of the first volume to occipital intralobar association fibers. The Dejerines observed five fiber bundles specific to the occipital lobe: the stratum calcarinum, the vertical occipital fasciculus (VOF) of Wernicke, the transverse bundle of the lingual lobule of Vialet, the transverse fasciculus of the cuneus and the stratum proprium cunei. At the time, intralobar white matter fibers were just being discovered – these were as yet poorly understood and surrounded by controversies due to inconsistent classification and nomenclature (Yeatman et al 2014; Mandonnet et al 2018), leading to much of the Dejerines’ work on their trajectory being overlooked.

Of late, the VOF, the largest of the occipital intralobar fibers dissected by the Dejerines, has been the focus of an increasing number of papers (Weiner et al 2017; Güngör et al 2017; Oishi et al 2018; Briggs et al 2018; Panesar et al 2019). As of December 2019, a PubMed search generated thirty hits relating to the “vertical occipital fasciculus” when used as a key term. In comparison, the other intraoccipital tracts generate no results and remain overlooked, mostly being listed in passing. An article by our group revisited the Dejerines’ work and provided access to the first English translation of the portion of the Dejerines’ book related to the long-range association fibers of the human brain (Bajada et al 2017). Here, we provide the first complete translation of the description by the Dejerines of the occipital originating short-range fibers trajectories, from the original French taken from *Anatomie des Centres Nerveux* (1895 p.780-784) (supplementary material 1 and 2). The aim of this study is to use the observations made by the Dejerines for guidance through a virtual dissection of the intralobar tracts of the occipital lobe, based on *in vivo* diffusion imaging data. The virtual dissections enable us, to have a better discern the three-dimensional relations of these once forgotten tracts.

## MATERIAL AND METHOD

The study consisted of two principal methods: the first being the translation from French into English of the Dejerines’ classical study of occipital intralobar fibers. This was followed by a diffusion weighted MRI (dMRI) based virtual reconstruction of the Dejerines’ dissection in healthy subjects.

### TRANSLATION

The masterful description of intra-occipital fiber tracts in Chapter 5: pp.780-4 by J.J. Dejerine and A. Dejerine-Klumpke in *Anatomie Centre Nerveux; Volume I* (1895) was fully translated (MB) from its original 19^th^-century French to more universally accessible English and reviewed for its accuracy by the experts in this field (CB, MTS). The complete translation is available as supplementary material.

### SUBJECT SELECTION

A template-based, probabilistic, fiber tractography study was conducted in 24 healthy and unrelated consenting volunteers (range 22-35 years, 12 males). The analysed diffusion MRI dataset was derived from the S1200 release of the publicly available WU-Minn Human Connectome Project (HCP) database (https://db.humanconnectome.org/). To ensure data homogeneity, the subjects selected had none of the following features/criteria: endocrine abnormalities including hyper and hypothyroidism, a handedness score below zero (people with a left-handed tendency), color vision abnormalities, illegal drug use, history of psychiatric problems, hypertensive individuals, alcohol detected by a breathalyser, data with quality control (QC) issues A or B. This study uses data collected and processed by the HCP and approved by the Washington University IRB. The additional data analysis conducted in this present study was approved by the local ethics committee of the University of Malta (UREC).

### IMAGE ACQUISITION

The full details of image acquisition can be found in the following articles (Moeller et al 2010; Feinberg et al 2010; Xu et al 2012; Setsompop et al 2012; Sotiropoulos et al 2013). In short, the HCP group generated the *in vivo* T_1w_ and diffusion weighted magnetic resonance imaging (MRI) of the human brains through a customised high-resolution 3-Tesla Siemens Skyra scanner equipped with a 32-channel head coil. Structural imaging utilised an axial 3D MPRAGE pulse sequence, while the DWI used a monopolar Sejskal-Tanner sequence (Van Essen et al 2012; Glasser et al 2013). Following the algorithm in Milchenko et al. (2013) DICOM image files were defaced and de-identified. For further data analysis, dcm2nii MRIcron was used to convert DICOM files to (NITRC)(http://www.nitrc.org/projects/mricron).

Diffusion data were rectified for head motion and geometrical distortion, and pre-processed through the HCP minimally processed pipeline (Jenkinson et al 2002; Andersson et al 2003; Fischl 2012; Jenkinson et al 2012; Glasser et al 2013; Andersson and Sotiropoulos 2015; Sampson et al 2015).

### MRI DATA PROCESSING

All additional processing was accomplished with the fiber tractography MRTrix3 (RC_3) software toolkit and iFOD2 algorithm (Tournier et al 2012; Tournier et al 2019). The fiber orientation distribution (FOD) functions from the diffusion signal in each voxel were computed using the multi-shell-multi-tissue constrained spherical deconvolution (CSD).

The individual FODs across the 24 subjects were used to create a population template within MRtrix3. A manually defined occipital lobe mask on the template was then used to produce occipital lobe specific connectomes. The safety margins between the occipital lobe and the exclusion border were large and included posterior aspects of the temporal and parietal lobe. The mask excluded the brainstem, the cerebellum. The anterior border of the mask was defined by a coronal line at the anterior aspect of the splenium of the corpus callosum that extended straight to the lateral convexity of the cerebrum. Figures of the mask are provided in supplementary material 3. The connectome derived from 10 million randomly seeded fibers from within it (iFOD2, Lmax= 8, Length: 10-150mm, Max angle = 45 degrees, output step size = 0.625mm, FOD cut-off = 0.05, FOD power = 3) (Tournier et al 2007; Jeurissen et al 2014). Fibers traversing beyond the occipital lobe were manually assessed. Inclusion and exclusion masks are available in the supplementary material.

## RESULTS

Using CSD tractography, this study successfully identified and mapped four of the five occipital intralobar white matter tracts observed by the Dejerines. The translation of the Dejerines’ work on these fibers, from 19^th^-century French to accessible English (Supplementary Material 1), helped guide the tractography findings. Tractography reconstruction of the occipital intralobar tracts have been overlaid on a 3D T_1w_ image and presented in figures 2-5. The relation of each tract to each other and to the longer association, projection and commissural fibers are provided in figure 6.

### OCCIPITAL DIFFUSION TRACTOGRAPHY

With regard to short-range association fibers arising from the occipital lobe, the Dejerines described the vertical occipital fasciculus (VOF) which spans the entirety of the occipital lobe lateral to the posterior ventricle, and four shorter association fiber tracts projecting superiorly and inferiorly from the banks of the calcarine fissure: the stratum calcarinum (SC), stratum proprium cunei (SPC), transverse fasciculus of the cuneus (TFC) and the transverse fasciculus of the lingual lobe of Vialet (TFV) (Fig 1).

**Figure 1.**
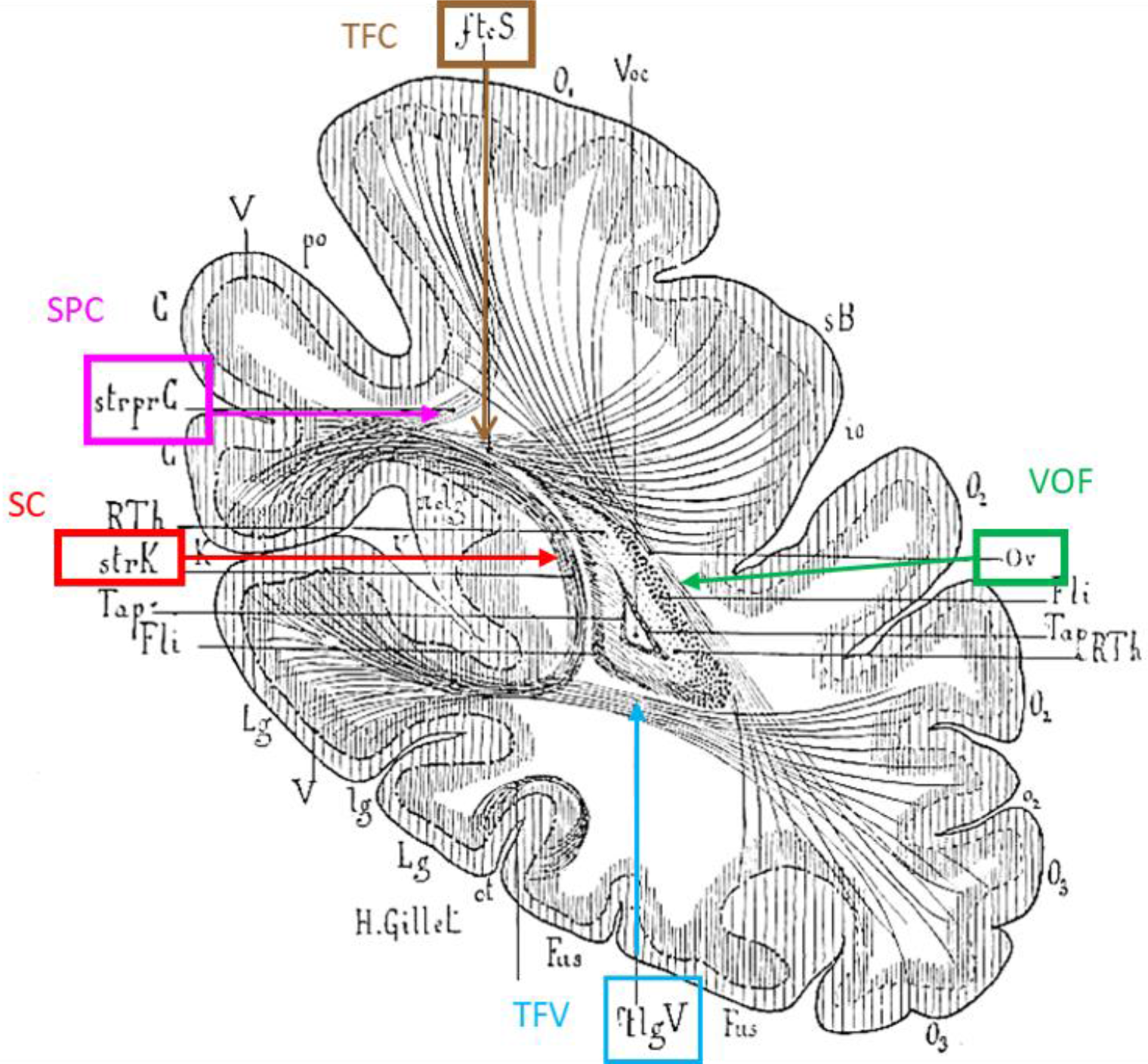
The Dejerines’ intraoccipital fibers. A coronal slice of the occipital lobe illustrated by H. Gillet depicting the occipital intralobar fibers identified by the Dejerines. VOF: vertical occipital fasciculus; TFV: transverse fasciculus of the lingual lobe of Vialet, SC: stratum calcarinum, SPC: stratum proprium cunei, TFC: transverse fasciculus of the cunei. Image adapted from Dejerine & Dejerine 1895, p. 781.

### STRATUM CALCARINUM (SC)

The stratum calcarinum (SC) consists of several U-shaped fibers that connect the two banks of the calcarine fissure, a major landmark that medially divides each occipital lobe into the wedge-shaped cuneus superiorly, and the lingual lobule inferiorly, and represents the boundaries of the primary visual cortex. This group of U-shaped fibers extends all along the calcarine fissure from the occipital pole to its junction with the parieto-occipital fissure at anterior border of the occipital lobe (Fig. 2). The SC is an unusually large U-fiber system because it directly connects adjacent gyri across a long and deep fissure – perhaps the reason the Dejerines devoted considerable attention to it, describing it alongside other intralobar fibers.

**Figure 2.**
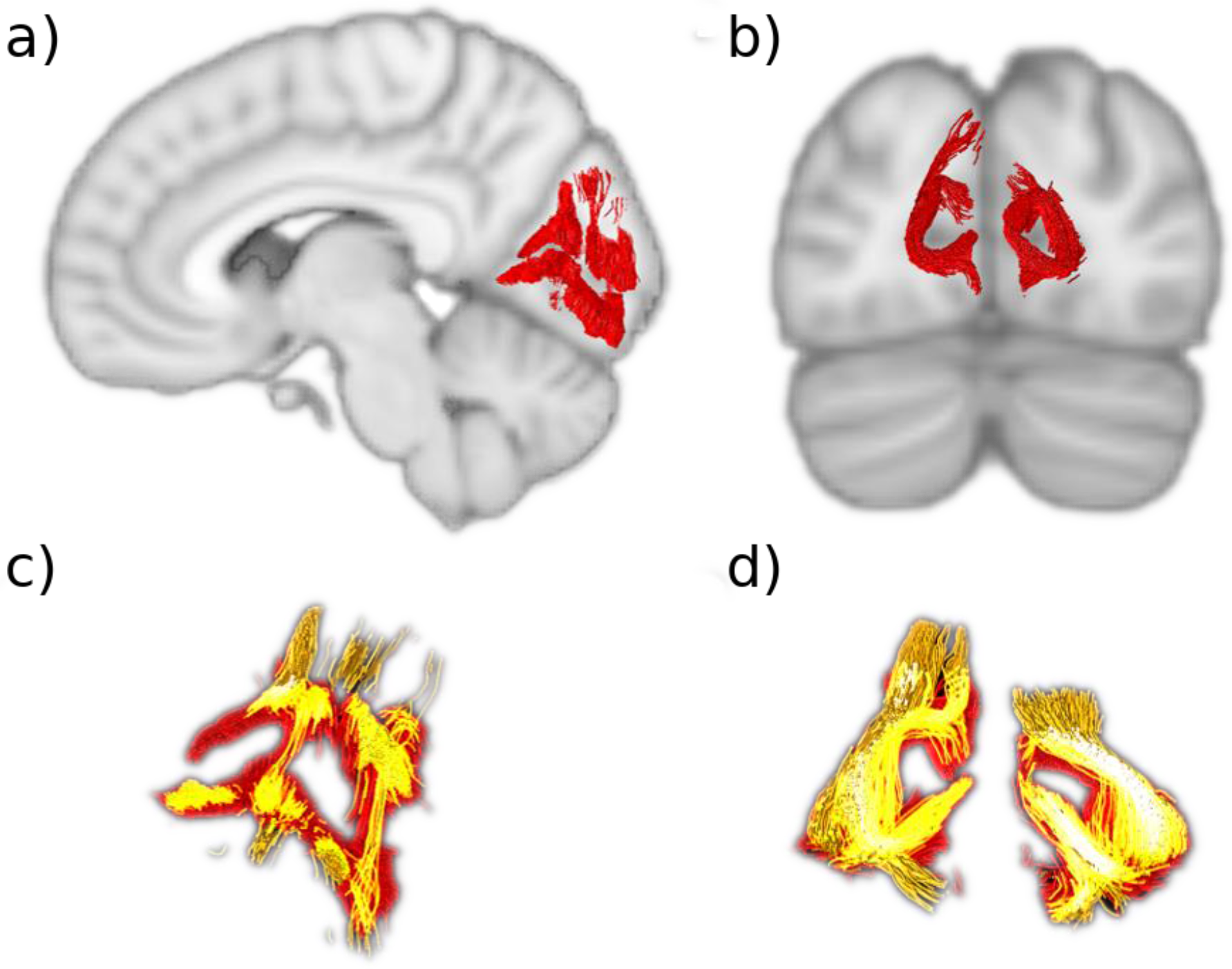
The stratum calcarinum (SC) A) Sagittal slice showing the right SC (*red*) curved around the calcarine fissure. B) Coronal slice showing the left and right SC. C-D) The two layers of the SC. The deeper and longer layer (*orange*) lies under the more superficial and u-shaped layer (*red*). Where they intersect is shown in *bright yellow*.

The SC fibers are composed of two superimposed layers (Fig. 2C). The first, most superficial layer has a vertical and horizontal orientation that embraces the superior and inferior edges of the fissure. The second has longer and more deeply penetrating fibers that join the medial-central aspects of the cuneus to the medial-central aspects of the lingual gyrus. SC layers lie directly on top of the longitudinal fibers of the inferior longitudinal fasciculus that joins them to carpet the floor of the calcarine fissure (Fig.6). These fibers run perpendicular to the vertical SC fibers and interlink the different U-fiber units along the fissure like rails on a track. This relation between the inferior longitudinal fasciculus and the U-fibers of the occipital lobe was depicted by ffytche et al. (2005), however, they did not explicitly mention the SC system or distinguish it from other U-fibers.

The horizontal aspect of the SC that projects away from the midline into the bank of the fissure becomes intertwined with the other intralobar tracts that originate from the lips of the calcarine fissure: the stratum proprium cunei (SPC in the superior lip) and the transverse fasciculus of Vialet (TFV in the inferior lip). In the anterior part of the fissure, the vertical portion of the SC is on the same plane, and slightly overlaps the sledge runner fibers (not described by the Dejerines) that travel vertically downwards, beneath the calcarine fissure, prior to continuing transversely to the anterior lingual gyrus (Fig. 6). Similarly, Meyer’s loop, the anterior optic radiation fiber bundle, terminates in the inferior lip amongst the SC.

### STRATUM PROPRIUM CUNEI (SPC)

The stratum proprium cunei (SPC) arises from the superior bank of the anterior calcarine fissure, just posterior to the junction with the parieto-occipital sulcus (Fig. 3). The tract connects the inferior aspect of the cuneus with the most anteromedial convexity of the occipital lobe, just posterior to the pre-cuneus. The SPC runs deep to the parieto-occipital sulcus, at the level of the calcar avis, initially coursing laterally away from the calcarine fissure for a short distance, before ascending vertically to terminate in what is classically referred to as the first occipital convolution. Unlike the vertical occipital fasciculus that fans out across the entirety of the first occipital convolution, the SPC projections remain on a plane medial to the posterior horn of the ventricle. Though the SPC connects distal regions of the same lobe, it is a relatively short and narrow association fiber tract. Our findings align with the Dejerines’, and show that the SCP runs neatly along the anteromedial margins but is constrained to the occipital lobe.

**Figure 3.**
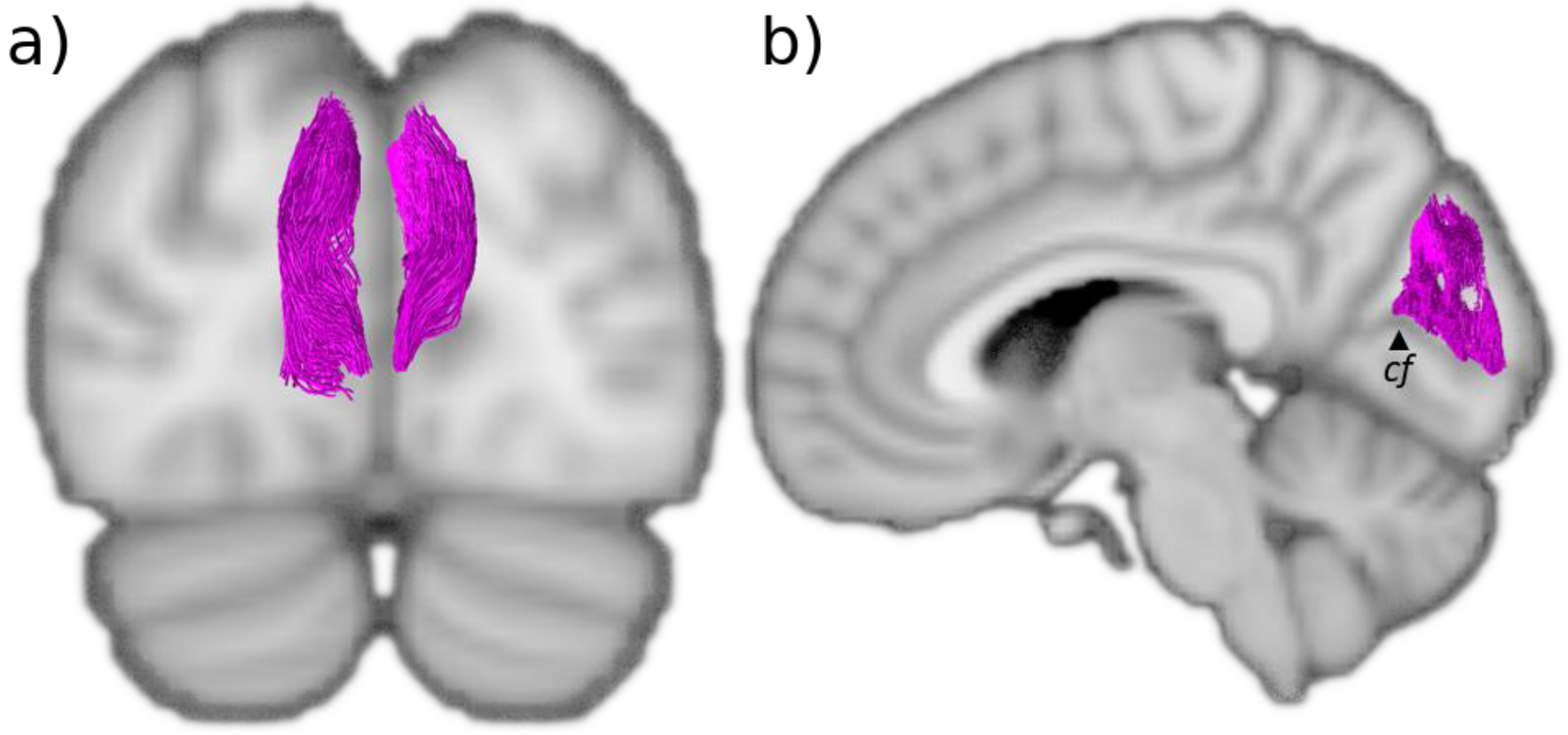
The stratum proprium cunei (SPC) A) Coronal view of the SPC (*magenta*) that lines the midline of the cuneus. B) Sagittal view of the SPC showing how it arises from the superior bank of the calcarine fissure. *cf*: calcarine fissure.

### TRANSVERSE FASCICULUS OF THE CUNEUS OF SACHS (TFC)

The transverse fasciculus of the cuneus of Sachs (TFC) is described as a tight, slender bundle, consisting of relatively few fibers that initially move away from the upper fold of the calcarine fissure with a strong oblique trajectory, such that they can course laterally over the roof of the posterior lateral ventricle and cross the cuneus to terminate in the lateral convexities of the occipital lobe (Sachs and Wernicke 1892; Vialet 1893; Campbell 1905; Greenblatt 1973; Schmahmann and Pandya 2006; Vergani et al 2014). Unfortunately, in the present study, it was not possible to identify a bundle resembling the TFC.

### TRANSVERSE FASCICULUS OF THE LINGUAL LOBE OF VIALET (TFV)

A tract resembling the many aspects of the Dejerines’ description of the transverse fasciculus of the lingual lobe of Vialet (TFV) was successfully isolated (Fig.4). The Dejerines depicted the TFV as a mirror counterpart to the transverse fasciculus of the cuneus, as it connects the inferior gyrus of the calcarine fissure to the infero-lateral aspects of the occipital lobe, akin to the way in which the transverse fasciculus of the cuneus connects the fissure to the external surface of the cuneus. In contrast, the stratum proprium cunei connects the fissure to the medial surface of the hemisphere and does not appear to have an inferior lingual gyrus equivalent. Interestingly, the TFV origin can be found all along the calcarine fissure alongside the stratum proprium cunei.

The TFV has a morphology that resembles an extended U-fiber; the tract travels laterally away from the fissure prior to adopting an intense anterior-oblique trajectory that sharply curves back on itself and continues in a posterior-oblique course toward the lateral convexity of the occipital lobe. Due to the squatness of the lingual lobe, the tract appears quite flat and two ends of the fiber tract are nearly on the same coronal plane, with the lateral terminations slightly inferior to its calcarine fissure origin. The TFV starts superiorly and medially to the inferior longitudinal fasciculus and occipital projections of the inferior fronto-occipital fasciculus, and crosses laterally over these two major association fiber bundles, before descending just laterally to them.

### VERTICAL OCCIPITAL FASCICULUS (VOF)

Using the Dejerines’ classical description, we identified the vertical occipital fasciculus (VOF) (Fig.5A), and it aligns well with modern tractography findings. Of the five occipital tracts the Dejerines observed, the VOF is the longest and most prominent, and consequently has been the focus of many more studies. Often described as a short, vertical, and ‘sheet-like’ bundle of fibers, the VOF “connects the superior aspect of occipital lobe to its inferior surface” (Déjerine and Déjerine-Klumpke 1895) p.779 maintaining a dorsal-ventral course lateral to the posterior ventricle. More specifically, the inferior terminations of the VOF extend from the ventral temporo-occipital cortex region, encompassing the inferior temporal gyrus and its intersection with the anterior aspects of the middle and inferior occipital gyri. The superior projections of the VOF reach the lateral gyri of the occipital lobe, including the transverse occipital sulcus, with several fibers reaching the posterior end of the intra-parietal sulcus and the angular gyrus.

**Figure 4.**
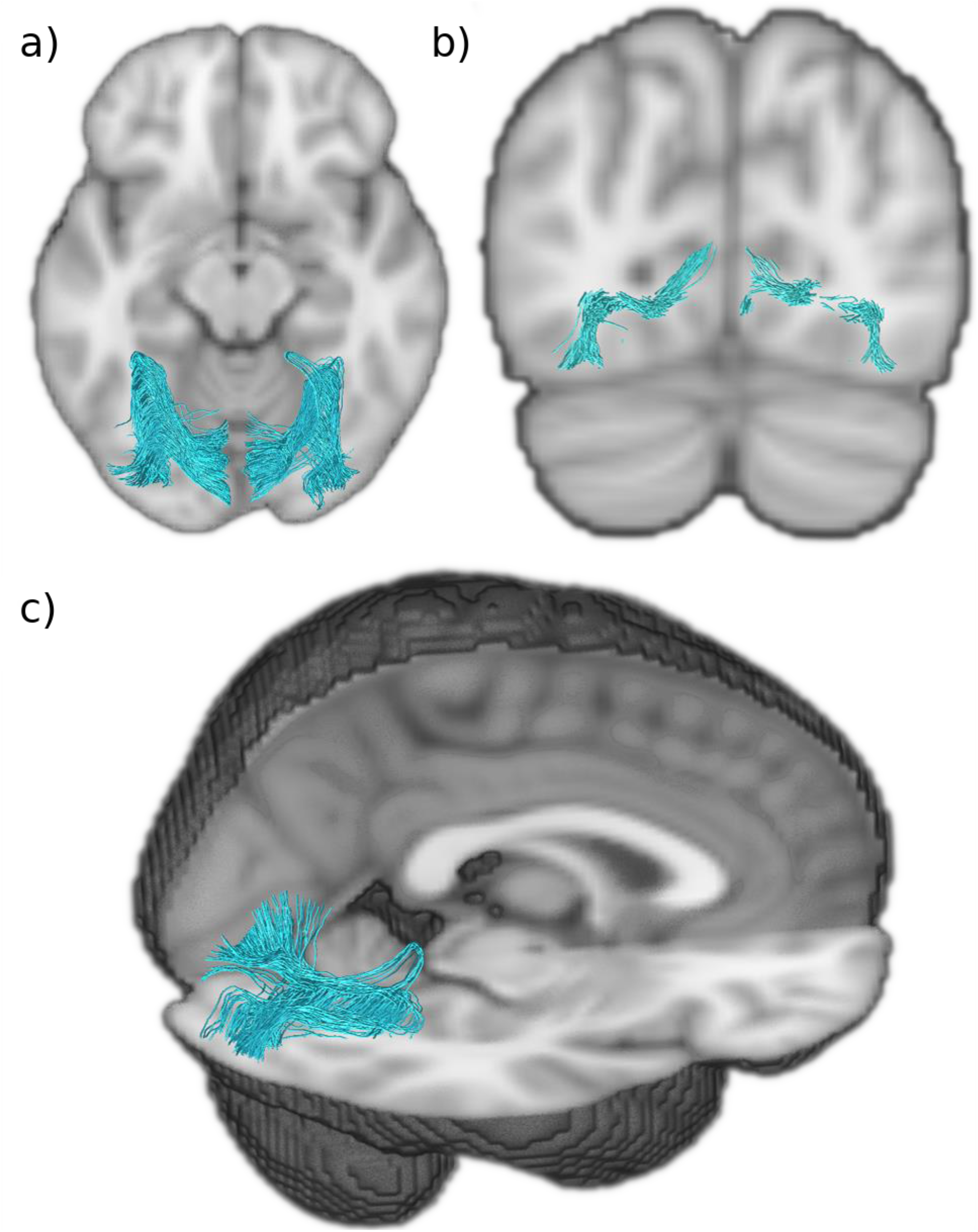
Transverse fasciculus of the lingual lobe of Vialet (TFV) A) Axial view of the TFV (*blue*) showing it projects anteriorly before it reflects in the anterior aspect of the lingual gyrus. B) Coronal view of TFV showing it arises from the inferior gyri of the calcarine fissure and terminates infero-laterally. C) Sagittal and axial view of the TFV showing how it does a hairpin loop at the anterior edge of the lingual gyrus.

**Figure 5.**
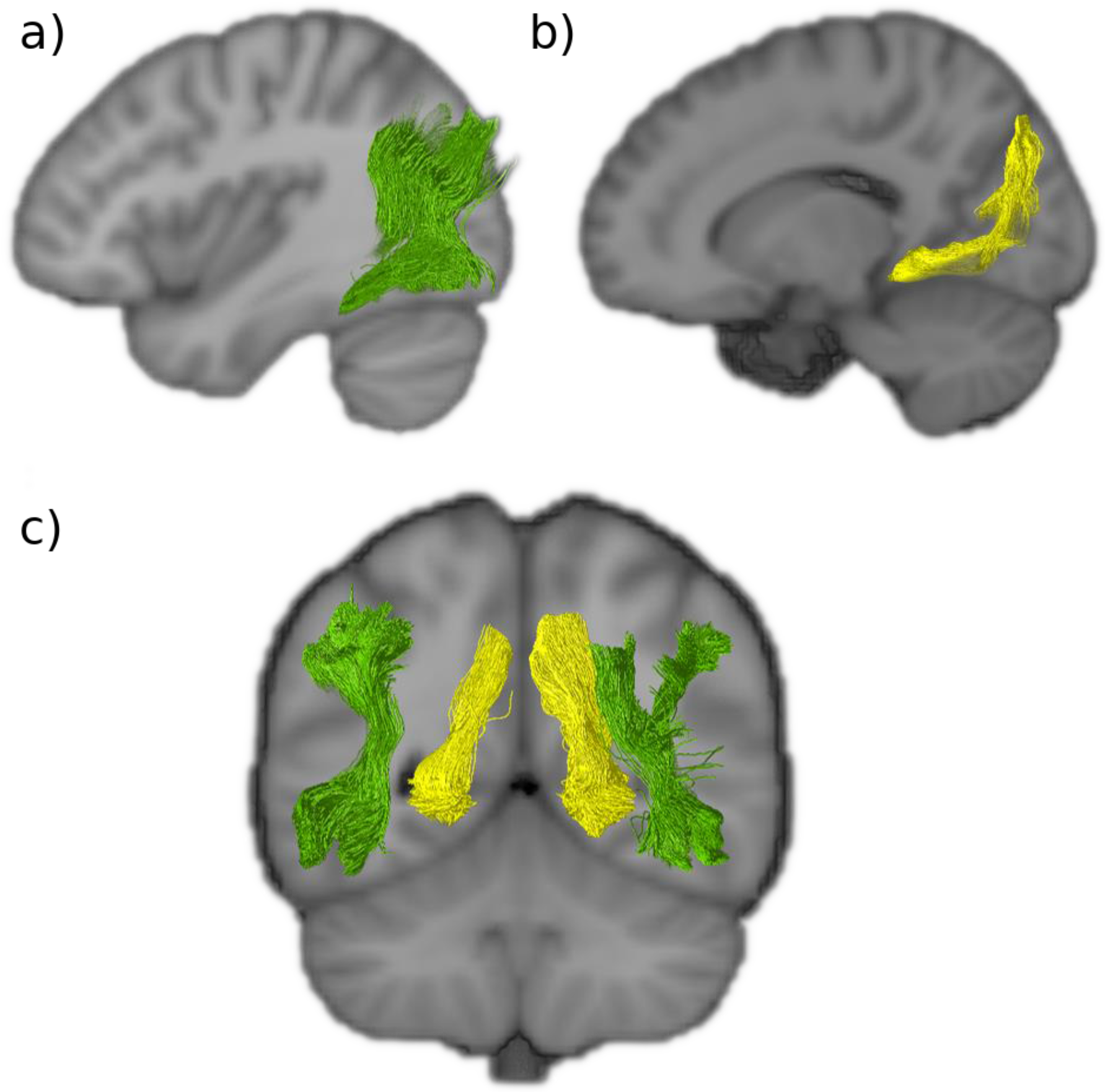
Dorsal to ventral intralobar fibers. A) Sagittal view of the right vertical occipital fasciculus (VOF in *green*). B) Sagittal view of the right sledge runner fasciculus (SFR in *yellow*). C) Coronal slice of the VOF and SFR showing how they lie lateral and medial to the lateral ventricles, respectively. Both connect dorsal aspects of the occipital lobe to ventral aspects. VOF: vertical occipital fasciculus, SFR: sledge runner fasciculus.

**Figure 6.**
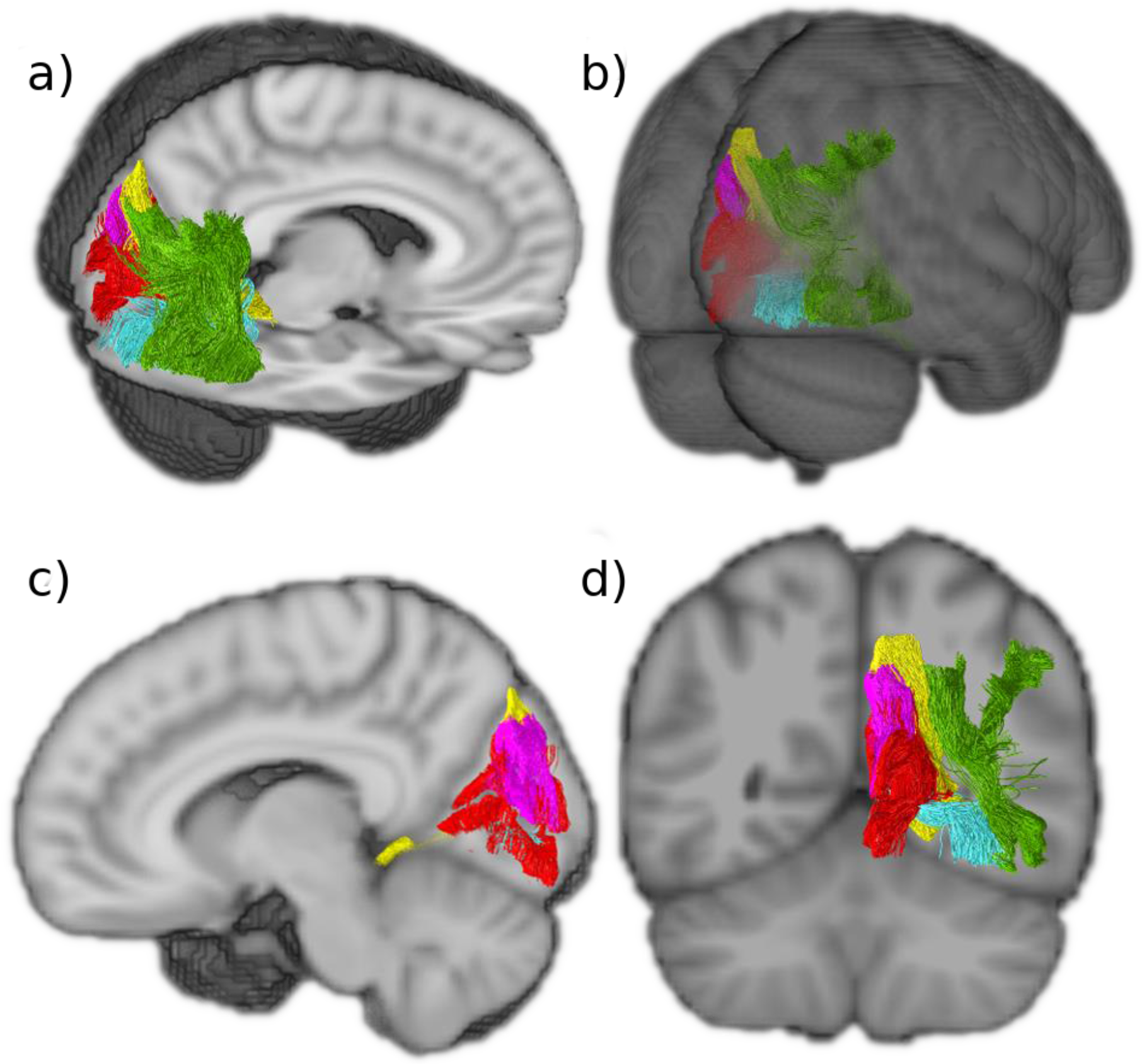
The five intralobar fibers identified in the occipital lobe. A) Sagittal and axial view. B) Posterior view. C) Sagittal view showing that the SPC is more medial than the SRF and that the SC extends more widely than the other medial intralobar tracts. D) Coronal view showing that the intralobar fibers connect all aspects of the occipital lobe VOF: vertical occipital fasciculus (*green*), SFR: sledge runner fasciculus (*yellow*), SCP: stratum proprium cunei (*magenta*), SC: stratum calcarinum (*red*), TFV: transverse fasciculus of the lingual lobe of Vialet (*blue*).

The ‘sheet-like’ stem of the VOF tract, that courses through the central part of the occipital lobe, is a wide bundle that has a predominantly vertical trajectory, apart from the ventral and dorsal terminations that flick out laterally and transversely to reach the convexity of the lateral occipital lobe across a wide area. Whereas the stratum calcarinum is covered laterally by the inferior longitudinal fasciculus, the VOF passes laterally to the inferior longitudinal fasciculus along with the inferior fronto-occipital fasciculus, continuing superiorly to both and coursing vertically, posterior to the arcuate fasciculus.

### SLEDGE RUNNER FASCICULUS (SRF)

Tracts resembling the recently discovered sledge runner fasciculus (SRF) (Vergani et al. 2014) were identified in both hemispheres and reconstructed successfully (Fig.5B). The SRF connects the posterior pre-cuneus to the anterior lingual gyrus of the lingual lobe, taking a strongly inferior anterior oblique trajectory with a couple of convexities – giving it a sledge morphology. It is a long occipital tract that traverses from the dorsal to ventral aspects of the cerebrum like the vertical occipital fasciculus. However, unlike the vertical occipital fasciculus, it is medial to the ventricles. With a compact central stalk that widens as the tract descends anteriorly and narrows superiorly, it is a prominent bundle of similar length to the vertical occipital fasciculus. As in the first observations of the SRF obtained through fiber microdissections, the results showed that the dorsal SRF terminations where located at the anterior border of the cuneus, adjacent to the midline of the cerebrum. The dorsal SRF terminations are entangled with the anterior fibers of the stratum proprium cunei bundle (Fig.5C). The ventral terminations are more anteriorly located and project deep into the lingual gyrus, posterior to the cingulate isthmus and the posterior part of the parahippocampal gyrus.

## DISCUSSION

The anatomical described in the present study suggests that the occipital lobe has at least five intrinsic white matter structures that fall between the long association fibers and the very short U-shaped fibers. These are: the stratum calcarinum, the transverse fasciculus of the lingual lobule of Vialet, the stratum proprium cunei, the vertical occipital fasciculus of Wernicke (VOF) – which were described by the Dejerines, and the recently discovered sledge runner fasciculus (SRF). Apart from the vertical occipital fasciculus of Wernicke and the sledge runner fasciculus, none have been the focus of dedicated studies since the turn of the 20^th^ century.

This current study also highlights the persistent utility of historical studies for guidance of neuroimaging findings.

### VERTICAL OCCIPITAL FASCICULUS (VOF)

The VOF’s existence as an association fiber bundle has been extensively verified, through dMRI studies (Yeatman et al 2014; Keser et al 2016; Briggs et al 2018; Panesar et al 2019; Schurr et al 2019), post-mortem dissections (Vergani et al 2014; Güngör et al 2017; Palejwala et al 2019), and it has even been identified and compared in macaques (Takemura et al 2016). Yet, the boundaries of the pathway remain unclear and disputed (Bullock et al 2019). Its vertical trajectory, like that of the stratum proprium cunei, distinguishes it from the other long association fibers that project into the occipital lobe, and its long dorso-ventral trajectory, located lateral to the ventricle, sets it apart from the intraoccipital tracts. The Dejerines, like other early neuroanatomists, admitted to poorly defining the anterior limits of the VOF. Their descriptions suggest that it extended anteriorly beyond the modern boundaries of the occipital lobe, descriptions borne out by most recent tractography studies, including this present one. Hence, it is arguable that the VOF is not strictly an autochthonous tract restricted to the occipital lobe (Weiner et al., 2017). Depending on how anterior the ventral terminations of the VOF were, it assumed either an oblique dorsal and posterior trajectory, or in the case of more posterior ventral terminations, a vertical trajectory to the occipital cortex. We did not find strong ventral VOF termination within the ventromedial-fusiform gyrus or the dorsomedial portion of the occipital lobe. Our findings show that the VOF extends from the dorsal to ventral aspects of the lateral parietal and occipital lobe as a thin sheet and terminates at the lateral-posterior end of the temporo-occipital region. As described by the Dejerines, the VOF courses laterally to the occipital horn, the inferior fronto-occipital fasciculus and inferior longitudinal fasciculus.

Studies now aim to better differentiate the VOF fibers from the neighbouring larger bundles. The VOF projects over the inferior longitudinal fasciculus. Their proximity once made the VOF difficult to distinguish from the terminal portions of the inferior longitudinal fasciculus, leading to its misidentification in one of the early studies on the macaque (Schmahmann and Pandya 2006). Similarly, the posterior arcuate fasciculus borders the anterior dorsal terminations of the VOF, and the two tracts can be joint or slightly separated depending on the subject (Curran 1909; Weiner et al 2017). Borders for the VOF, or any other occipital intralobar tract, can manifest small discrepancies across studies partly due to brain borders being a man-made construct, high subject variability, the limits of imaging resolution, and the larger number of more prominent tracts that cross the intralobar tracts (Maier-Hein et al 2017). The occipital intralobar tracts, notably the VOF, have a strong dorsal-ventral trajectory, whereas many of the major long association fiber pathways that enter the occipital lobe have an anterior-posterior trajectory. Hence, the intralobar fibers cross paths with more prominent bundles that can cancel out their signal. The Dejerines themselves remarked on the limits of even masterful brain dissection skills for identification of terminations in pathway-dense regions, limiting the extent of guidance that post-mortem dissections can provide. The continuous improvement in MRI tractography angular resolution, and the development of alternative models, have made some headway (Takemura et al 2016; Weiner et al 2017). However, the extent of the separation between adjacent fiber pathways is subject to the imaging modality employed (Weiner et al 2017). It has been shown that the VOF and the posterior arcuate fasciculus are seen to have closer borders when using constrained spherical deconvolution based probabilistic tractography, compared to tensor based deterministic tractography (Weiner et al 2017). Recent studies have differentiated the VOF bundle from the vertical subdivision of the arcuate fasciculus (Panesar et al. 2019). It is important to make these distinctions, for both the VOF and other intralobar fibers, in order to generate a truly representative map – that is, one that does not include pathways that appear longer or wider than they truly are because they artificially contain subsections of adjacent tracts. Knowing the extent of a distinct white matter system is useful for future work concerning functional specialization and pathology. It has been suggested that certain U-fibers, situated between the VOF and other long association pathways, form artificial connections, or U-fiber ‘bridges’, between the pathways that can make the two tracts look like one larger tract (Panesar et al 2019). The intermingling of U-fibers is known to obstruct the visualisation and interpretation of the deeper underlying subcortical tracts in both fiber tractography and dissections (Reveley et al 2015).

The difficulty that arises when studying the work of the Dejerines, like other 19^th^-century anatomists, relates to translating their definition of structures into the current school of thought. Inconsistent landmarks, nomenclatures and methods were used to define and visualise structures. This discordance makes it difficult to reference a classic text and persists to the present day. Nevertheless, the congruency between the Dejerines’ dissection and modern imaging suggests that the other tracts they described should also be successfully visualised in vivo.

### STRATUM CALCARINUM (SC)

In vivo and in vitro dissections of the calcarine fissure reveal a large continuous bundle of U-fibers, named the stratum calcarinum (SC), connecting the upper and lower edges of calcarine cortex (Fig.2). This is the first tractography reconstruction of the SC as U-fiber studies are infrequent. U-fibers typically do not form a clear pathway or large fiber system. The Dejerines chose to omit the SC from the portion of their book that concerns U-fibers, in favour of discussing it alongside the association fibers of the occipital lobe to highlight the prominence and size of the SC. The Dejerines emphasised that it is not a true association fiber or a single entity. Instead, it represents a series of tracts forming an oversized U-fiber system that curves around the whole calcarine fissure and is specific to the primary visual cortex. It extends from the posterior end of the occipital horn to the intersection with parieto-occipital fissure and circumnavigates the calcar avis in its full extension. The SC intermingles with the other association fibers like the transverse fasciculus of Vialet and stratum proprium cunei, which link the primary visual cortex with the convexity of the occipital lobe.

The SC system forms an atypical bundle as it consists of a long and deep layer underlying a short and more superficial layer. The superficial SC layer, as can be seen from the results (fig. 2C), has short projections that tightly carpet the floor of the calcarine fissure and join the superior and inferior gyri that directly border the fissure. This gives the layer the characteristic ‘U’ appearance. The second underlying layer has wider projections and a weaker U shape. Its longer fibers connect the medial aspect of the central cuneus to the inferior-medial aspect of the lingual lobe.

The calcarine fissure receives several white matter bundle pathways that share similar orientations at the level of the fissure, making it particularly difficult to differentiate them. Campbell (1905), postulated that the majority of the fibers labelled by the Dejerines, such as the SC, were actually terminal fibers of the optic radiation. Indeed, the Dejerines (Déjerine and Déjerine-Klumpke, 1895) acknowledged that the histological staining method employed was insufficient to confidently evaluate the degree to which association fibers, like the SC, contribute to the projection fiber pathways that are situated along the ventricles. However, more recent studies, conducted with the improved Klingler’s technique, are congruent with the Dejerines’ description of the SC (Vergani et al 2014; Koutsarnakis et al 2019). Unfortunately, to date, most mentions of the human SC, transverse fasciculus of Vialet, transverse fasciculus of the cuneus, and stratum proprium cunei in the literature have been in reference to historical dissections (Catani and De Schotten 2012; Catani et al 2012; Takemura et al 2018).

A human post-mortem dissection study had shown the three-dimensional relationship between the sledge runner fasciculus and SC, with the sledge runner fasciculus running obliquely, almost perpendicularly, over the SC fibers that lie in the anterior portion of the calcarine fissure just posterior to the junction it shares with the parieto-occipital sulcus (Koutsarnakis et al 2019). There is a similar overlap that occurs between the SC and the stratum proprium cunei (fig.6). The posterior inferior longitudinal fasciculus projects obliquely toward the calcarine fissure and passes over the SC, before descending and coating the internal wall of the occipital horn, separating the ventricle from the SC (Déjerine and Déjerine-Klumpke 1895; Schmahmann and Pandya 2006).

The Dejerines considered the SC as one of the most important short-range connections in the occipital lobe. The proposed importance of the SC arises from its role in connecting the superior and inferior vision field regions all along the fissure, likely implicating it in the visual pathways of the macula and the retina. It has been postulated that the SC, stratum proprium cunei and transverse fasciculus of the cuneus contribute to coordination and propagation of the visual stimuli of colour, shape and motion from the primary cortex to higher order association areas, permitting both dorsal and ventral outflow to the rest of the parietal and temporal cortex (Greenblatt 1973; Maunsell and van Essen 1983).

### STRATUM PROPRIUM CUNEI (SPC)

The SPC is a short and seemingly inconsequential tract that rises in the sagittal plane from the calcarine fissure at the medial border of the hemisphere (fig.3). Since the first reports by Sachs (Sachs and Wernicke 1892), it has only been vaguely mentioned in passing (Campbell 1905; Von Bonin et al 1942; Greenblatt 1973; Schmahmann and Pandya 2006; Vergani et al 2014). Our current findings align strongly with the Dejerines’ observation that the SPC “originates from the superior lip of the calcarine fissure” and projects dorsally to the “cortical region of the superior border of the hemisphere” (Déjerine and Déjerine-Klumpke 1895), p.782). The results show a bundle with a clear vertical projection that links the upper border of the anterior calcarine fissure to the medial aspect of the cuneus convexity. Like the stratum calcarinum, the SPC extends along the entire length of the calcarine fissure. It differs, however, by being limited to the cuneus. Schmahmann & Pandya (2006), speculated that the SPC bundle corresponds to fibers they identified in rhesus macaque, located in the superior parietal lobule and the dorsal occipital white matter, caudal to the dorsal component of the superior longitudinal fasciculus. The fibers connected the medial pre-occipital gyrus of the occipital lobe with the medial part of the superior parietal lobule. However, the SPC we observed does not extend past the parieto-occipital fissure and does not correspond with the observations of Schmahmann & Pandya, 2006, who describe it as more anteriorly projecting compared to the system we have identified. As this is the first reconstruction of the SPC with diffusion tractography, it is not clear to what extent the approach’s limitations have affected the reconstruction.

### TRANSVERSE FASCICULUS OF THE LINGUAL LOBE OF VIALET (TFV)

The Dejerines named the TFV after Nehemie Vialet who described it in detail and published a paper arguing for its existence as a distinct fascicle (Vialet 1893). The Dejerines were able to successfully reproduce the observations made by Vialet in their own dissection studies. Our study succeeded in reconstructing this bundle (fig.4). This was unexpected, as like the transverse fasciculus of the cuneus, the TFV tract has not been mentioned in recent occipital fiber dissection or tractography studies. The TFV tract can be seen as a counterpart to the transverse fasciculus of the cuneus, as it connects the inferior gyri of the calcarine fissure to the convexity of the lingual lobule. The TFV can be differentiated from the inferior longitudinal fasciculus that courses alongside it at the level of the lingual gyrus and posterior temporal lobe, by its slightly thinner bulk and paler Pal-haematoxylin staining (Vialet 1893; Déjerine and Déjerine-Klumpke 1895). Vialet considered this bundle to be functionally important as it seems to represent the inferior portion of the occipital lobe association fiber system.

The existence of the TFV, as a distinct entity alongside the transverse fasciculus of the cuneus and vertical occipital fasciculus, was heavily contested shortly after their discoveries. This resulted in their omission from much of the literature, such as the neuroatlases of notable and influential neurologists like that of Constantin Von Monakow. Von Monakow described various subcortical structures with the locations and gross trajectories that match the TFV, transverse fasciculus of the cuneus and vertical occipital fasciculus, but was unable to differentiate them from neighbouring bundles and so negated their existence as distinct tracts (Yeatman et al 2014). Indeed, Schmahman & Pandya (2006), professed uncertainty as to the existence of the TFV as a distinct tract and considered the TFV as described by the Dejerines to be equivalent to the *“transverse inferior longitudinal fasciculus fibers that lie within the ventral part of the occipital and temporal lobe and that link the lateral cortices with the medial cortices”* (p. 450). However, Schmahmann & Pandya, 2006, had made a similar comment in relation to the vertical occipital fasciculus and had interpreted it as part of the vertical component of the inferior longitudinal fasciculus (Forkel et al 2014), which was later shown to be inaccurate by more focused tractography studies. Vergani et al. (2014), conducted a thorough post-mortem exploration of the intralobar fibers of the occipital lobe, in an effort to replicate the work of Sachs (Sachs and Wernicke 1892). Interestingly, like Sachs, the authors did not mention a tract that matches the morphology of the TFV as described later by Vialet, the Dejerines, and as identified in this study. A recently resurfaced photograph of a Weigert-stained histological slide of the macaque occipital lobe by von Bonin et al. (von Bonin et al., 1942) clearly depicts the vertical occipital fasciculus and stratum calcarinum, and somewhat less clearly, the TFV and transverse fasciculus of the cuneus named the fasciculus transversus lingualis and fasciculus transversus cunei, respectively (Takemura et al., 2018). The brevity of the description makes it hard to ascertain in greater detail the course of the TFV in the macaque.

### THE TRANSVERSE FASCICULUS OF THE CUNEUS (TFC)

Our study did not identify a consistent tract in either hemisphere that matches any previous descriptions. Yet, the Dejerines remarked that the calcarine cortex is attached to the cuneus and first occipital convolution through both the stratum proprium cunei and TFC and this observation of the TFC was also made by (Sachs and Wernicke 1892; Vialet 1893; Campbell 1905; Von Bonin et al 1942; Greenblatt 1973; Vergani et al 2014). The Dejerines, with reference to Sachs’ work, described the TFC as having a trajectory that “runs slightly obliquely, anteriorly and laterally, and radiate[s] in the superior parietal lobule and in the angular gyrus” (Dejerine, 1895 pg.781). It was specified by the Dejerines, Sachs and Vialet (Sachs and Wernicke 1892; Vialet 1893; Déjerine and Déjerine-Klumpke 1895) that the origin of the TFC is at the level of the calcarine fissure amongst the stratum calcarinum and stratum proprium cunei. This was more recently supported by Vergani et al., 2014, (Vergani et al 2014) who conducted an in vitro dissection and identified segments of the TFC, which they labelled stratum transverse cunei. The cuneus is very compact with a high density of white matter fibers crossing it. Therefore, there are a large amount of fibers with an antero-posterior trajectory, crossing paths with the thinner TFC, either perpendicular or parallel to it, disrupting the fiber orientation. Indeed, a recent effort to follow the full trajectory of this bundle was unsuccessful (Vergani et al., 2014).

The two opposing origins along the calcarine fissure of the transverse fasciculus of Vialet and TFC, and the mirroring trajectories described by the Dejerines, imply the possibility of complementary functions related to the upper and lower quadrants of the visual field, respectively, and integration of visual stimuli to the accessory visual integration area and the rest of the cerebrum (Campbell 1905; Greenblatt 1973). It follows that if the transverse fasciculus of Vialet exists to connect the areas of the cortex involved in the superior vision fields with distal cortical regions, a similar subcortical system would need to be in place for the inferior vision fields.

### THE SLEDGE RUNNER FASCICULUS (SRF)

The SRF had not been described by Sachs, nor later by the Dejerines or other neuroanatomists until Vergani and colleagues (Vergani et al 2014). The novel tract was initially named the ‘sledge-runner fasciculus’ to highlight its undulating shape due to its anteriorly projecting curve at the level of the calcarine fissure (Vergani et al 2014). Later, in a focused tractography study, the SFR was alternatively named as the medial occipital longitudinal tract (MOLT) to better reflect its anatomy (Beyh et al 2017). We continue to use the term SRF as it is the more widely known term. The SRF is a thin tract that lies within the medial aspect of the anterior calcarine fissure relaying the anterior border of the dorsal occipital lobe to the antero-superior aspect of the lingual gyrus. Previous tractography studies align with our observation that the SRF lies deep to the anterior third of the stratum calcarinum. The SRF connects the anterior border of the medial cuneus to the lingula, isthmus of the cingulum and posterior parahippocampal gyrus (Vergani et al 2014; Güngör et al 2017; Koutsarnakis et al 2019). Vergani et al. (2014), supposed that previous anatomists failed to identify the SRF because they classically examined the brain using coronal sections. This allowed U-fibers, running in the coronal plane along the medial and inferior aspects of the hemisphere, to mask the tract. Subsequently, studies combining both post-mortem dissections and tractography confirmed the dorsomedial-ventrolateral trajectory of the SRF, specifying that it connects the posterior part of the precuneus to the lingual gyrus and that it lies predominantly medial to the occipital horn and the posterior two-thirds of the atrium (Güngör et al 2017; Koutsarnakis et al 2019). The SRF abuts the posterior cingulum and passes medial to the forceps major with a convex anterior projection.

Furthermore, based on current functional understanding of the regions near the SRF tract dorsal origins, specifically the posterior precuneus, it is postulated that the SRF has a role in processing visual stimuli for visuospatial attention and the recognition of places (Güngör et al 2017; Koutsarnakis et al 2019). The present study is in concordance with past findings and shows the SRF’s relations to other intraoccipital tracts (fig.6).

### LIMITATIONS

As we have shown throughout this study, dMRI has permitted the explorative and comparative studies of subcortical architecture and morphology in large pools of healthy patients, producing both qualitative and quantifiable *in vivo* data. Regardless, it is necessary to acknowledge the inherent limitations of dMRI based tractography and the specific shortcomings of the technique used in our study. First, it is well documented that the foundation of all successful white matter studies is a strong anatomical background to limit tract misinterpretation and erroneous definitions (Thomas et al 2014; Maier-Hein et al 2016). Interpretation of the results depends on the expertise of the operator in both physical and virtual dissections. Second, literature has extensively noted that tractography struggles to discern crossing and kissing fibers (Maier-Hein et al 2016). Third, the inaccurate localisation of tract endpoints is a persistent issue despite the improvement in image resolution (Smith et al 2012; Girard et al 2014). Identification of endpoints requires heuristic selection techniques and uses varied termination criteria, introducing a considerable operator bias. Fourth, selecting a method that prevents spurious inferences by limiting the trade-off between sensitivity and specificity, when reconstructing tracts an unsolved problem (Thomas et al 2014; Zalesky et al 2016). Furthermore, the anatomical accuracy of tractography relies on the selected algorithm parameters, with their optimisation varying from bundle to bundle. The choice of these parameters can lead to biased characterisation of connectome topography and therefore remains a source of controversy.

One of the standard approaches to try to reduce the spurious results of tractography, particularly false positives, is the implementation of anatomical knowledge into the interpretation of the results. A form of anatomical confirmation is invasive post-mortem human brain dissection. This study consulted the work of classical neuroanatomists, such as the Dejerines, to guide *in vivo* dissections.

With the growing sparsity of appropriate brain cadavers, the development of tractography has used the historic work of past neuroanatomists to supplement their imaging findings (Schmahmann and Pandya 2007; Schmahmann et al 2008; Catani et al 2012; Dick and Tremblay 2012; Bullock et al 2019). The 19^th^-century neuroanatomists were masterful in their blunt post-mortem dissections and histological staining studies. The Dejerines, and the other neuroanatomists they reviewed in their work, used brains that were not necessarily healthy, did not use the Klingler’s technique, nor was there a clear consensus on the definition of association fibers. The first two likely led to artefacts and unclear structures, and the latter incorporated a bias into the interpretation of their findings. There are therefore limits to using historical dissection work to guide new findings. This is especially the case since different neuroanatomists omitted certain tracts from their atlases that they judged, rightly or wrongly, to be inaccurate. Historical work has proven to be a useful starting point for reviving unexploited concepts. It also serves as a strong guideline in helping to determine the potential value of a study. Regardless of the above, they do not circumvent the need for subsequent validation from the modern post-mortem dissection literature.

Tractography has generated findings shrouded in controversies, many dating from the 19^th^ century that are still unresolved. Yet, great strides have been made in acknowledging and reducing biases. In this study we have shown that there is a significant concordance between the original anatomical work of the Dejerines and the recently well-defined VOF, giving us confidence to extrapolate trust in our findings of the other lesser-known tracts. The results indicate gaps in current knowledge and encourage future studies based on this new perspective of what the association fibers of the occipital lobe entail.

## ACKNOWLEDGEMENTS

This article is based upon work from COST Action CA18106, supported by COST (European Cooperation in Science and Technology). SC received funding via the Helmholtz Initiative and Networking Fund as well as from the European Union’s Horizon 2020 Research and Innovation Program under Grant Agreement 785907 (Human Brain Project SGA2) and 945539 (Human Brain Project SGA3). MTS is funded by the European Research Council (ERC) under the European Union’s Horizon 2020 research and innovation programme (Grant agreement No. 818521).

The authors would like to thank Krista Bonnello Rutter Giappone for her meticulous critique on language and style of this manuscript.

Data were provided by the Human Connectome Project, WU-Minn Consortium (Principal Investigators: David Van Essen and Kamil Ugurbil; 1U54MH091657) funded by the 16 NIH Institutes and Centers that support the NIH Blueprint for Neuroscience Research; and by the McDonnell Center for Systems Neuroscience at Washington University.

## REFERENCES

Andersson JLR, Skare S, Ashburner J (2003) How to correct susceptibility distortions in spin-echo echo-planar images: application to diffusion tensor imaging. Neuroimage 20:870–888. doi: 10.1016/S1053-8119(03)00336-7

Andersson JLR, Sotiropoulos SN (2015) Non-parametric representation and prediction of single- and multi-shell diffusion-weighted MRI data using Gaussian processes. Neuroimage 122:166–176. doi: 10.1016/j.neuroimage.2015.07.067

Bajada CJ, Banks B, Lambon Ralph MA, Cloutman LL (2017) Reconnecting with Joseph and Augusta Dejerine: 100 years on. Brain 140:2752–2759. doi: 10.1093/brain/awx225

Bao Y, Wang Y, Wang W, Wang Y (2017) The Superior Fronto-Occipital Fasciculus in the Human Brain Revealed by Diffusion Spectrum Imaging Tractography: An Anatomical Reality or a Methodological Artifact? Front Neuroanat 11:119. doi: 10.3389/fnana.2017.00119

Beyh A, Laguna Luque P, De Santiago Requejo F, et al (2017) The medial occipital longitudinal tract: a white matter system for spatial navigation.

Bogousslavsky J (2005) The Klumpke family--memories by Doctor Déjerine, born Augusta Klumpke. Eur Neurol 53:113–120. doi: 10.1159/000085554

Briggs RG, Conner AK, Sali G, et al (2018) A Connectomic Atlas of the Human Cerebrum-Chapter 16: Tractographic Description of the Vertical Occipital Fasciculus. Oper Neurosurg (Hagerstown) 15:S456–S461. doi: 10.1093/ons/opy270

Bullock D, Takemura H, Caiafa CF, et al (2019) Associative white matter connecting the dorsal and ventral posterior human cortex. Brain Struct Funct 224:2631–2660. doi: 10.1007/s00429-019-01907-8

Campbell A (1905) Histological studies on the localisation of cerebral function. Cambridge University Press, Cambridge

Catani M, De Schotten MT (2012) Atlas of Human Brain Connections. Oxford University Press, Oxford

Catani M, Dell’acqua F, Vergani F, et al (2012) Short frontal lobe connections of the human brain. Cortex 48:273–291. doi: 10.1016/j.cortex.2011.12.001

Chou W, Salamon G, Orr NB, Salamon N (2009) Neuroanatomic analysis of diffusion tensor imaging of white matter tracts with dejerine sections and neuroimaging. Neuroradiol J 22:499–517. doi: 10.1177/197140090902200501

Curran EJ (1909) A new association fiber tract in the cerebrum with remarks on the fiber tract dissection method of studying the brain. J Comp Neurol Psychol 19:645–656. doi: 10.1002/cne.920190603

David S, Heemskerk AM, Corrivetti F, et al (2019) The superoanterior fasciculus (SAF): A novel white matter pathway in the human brain? Front Neuroanat 13:24. doi: 10.3389/fnana.2019.00024

Dejerine JJ, Dejerine-Klumpke A (1902) Anatomie des Centres Nerveux. Tome 2. Rueff et Cie, Paris

Déjerine JJ, Déjerine-Klumpke A (1895) Anatomie des centres nerveux: Tome 1. Rueff et Cie, Paris

Dick AS, Tremblay P (2012) Beyond the arcuate fasciculus: consensus and controversy in the connectional anatomy of language. Brain 135:3529–3550. doi: 10.1093/brain/aws222

Donahue CJ, Sotiropoulos SN, Jbabdi S, et al (2016) Using Diffusion Tractography to Predict Cortical Connection Strength and Distance: A Quantitative Comparison with Tracers in the Monkey. J Neurosci 36:6758–6770. doi: 10.1523/JNEUROSCI.0493-16.2016

Feinberg DA, Moeller S, Smith SM, et al (2010) Multiplexed echo planar imaging for sub-second whole brain FMRI and fast diffusion imaging. PLoS One 5:e15710. doi: 10.1371/journal.pone.0015710

ffytche DH, Catani M (2005) Beyond localization: from hodology to function. Philos Trans R Soc Lond B, Biol Sci 360:767–779. doi: 10.1098/rstb.2005.1621

Fischl B (2012) FreeSurfer. Neuroimage 62:774–781. doi: 10.1016/j.neuroimage.2012.01.021

Forkel SJ, Thiebaut de Schotten M, Kawadler JM, et al (2014) The anatomy of fronto-occipital connections from early blunt dissections to contemporary tractography. Cortex 56:73–84. doi: 10.1016/j.cortex.2012.09.005

Girard G, Whittingstall K, Deriche R, Descoteaux M (2014) Towards quantitative connectivity analysis: reducing tractography biases. Neuroimage 98:266–278. doi: 10.1016/j.neuroimage.2014.04.074

Glasser MF, Sotiropoulos SN, Wilson JA, et al (2013) The minimal preprocessing pipelines for the Human Connectome Project. Neuroimage 80:105–124. doi: 10.1016/j.neuroimage.2013.04.127

Greenblatt SH (1973) Alexia without agraphia or hemianopsia. Anatomical analysis of an autopsied case. Brain 96:307–316. doi: 10.1093/brain/96.2.307

Güngör A, Baydin S, Middlebrooks EH, et al (2017) The white matter tracts of the cerebrum in ventricular surgery and hydrocephalus. J Neurosurg 126:945–971. doi: 10.3171/2016.1.JNS152082

Hau J, Sarubbo S, Perchey G, et al (2016) Cortical Terminations of the Inferior Fronto-Occipital and Uncinate Fasciculi: Anatomical Stem-Based Virtual Dissection. Front Neuroanat 10:58. doi: 10.3389/fnana.2016.00058

Jbabdi S, Behrens TE (2013) Long-range connectomics. Ann N Y Acad Sci 1305:83–93. doi: 10.1111/nyas.12271

Jbabdi S, Sotiropoulos SN, Haber SN, et al (2015) Measuring macroscopic brain connections in vivo. Nat Neurosci 18:1546–1555. doi: 10.1038/nn.4134

Jenkinson M, Bannister P, Brady M, Smith S (2002) Improved optimization for the robust and accurate linear registration and motion correction of brain images. Neuroimage 17:825–841. doi: 10.1006/nimg.2002.1132

Jenkinson M, Beckmann CF, Behrens TEJ, et al (2012) FSL. Neuroimage 62:782–790. doi: 10.1016/j.neuroimage.2011.09.015

Jeurissen B, Tournier J-D, Dhollander T, et al (2014) Multi-tissue constrained spherical deconvolution for improved analysis of multi-shell diffusion MRI data. Neuroimage 103:411–426. doi: 10.1016/j.neuroimage.2014.07.061

Jeurissen et al. B (2017) Diffusion MRI fiber tractography of the brain. https://drive.google.com/file/d/14tAjvFdTBWwzH3payWFeGWIy4_FH5jwp/view. Accessed 7 Mar 2018

Kamali A, Flanders AE, Brody J, et al (2014) Tracing superior longitudinal fasciculus connectivity in the human brain using high resolution diffusion tensor tractography. Brain Struct Funct 219:269–281. doi: 10.1007/s00429-012-0498-y

Keser Z, Ucisik-Keser FE, Hasan KM (2016) Quantitative Mapping of Human Brain Vertical-Occipital Fasciculus. J Neuroimaging 26:188–193. doi: 10.1111/jon.12268

Koutsarnakis C, Kalyvas AV, Skandalakis GP, et al (2019) Sledge runner fasciculus: anatomic architecture and tractographic morphology. Brain Struct Funct 224:1–16. doi: 10.1007/s00429-018-01822-4

Krestel H, Annoni J-M, Jagella C (2013) White matter in aphasia: a historical review of the Dejerines’ studies. Brain Lang 127:526–532. doi: 10.1016/j.bandl.2013.05.019

Maier-Hein K, Neher P, Houde J-C, et al (2016) Tractography-based connectomes are dominated by false-positive connections. BioRxiv. doi: 10.1101/084137

Maier-Hein KH, Neher PF, Houde J-C, et al (2017) The challenge of mapping the human connectome based on diffusion tractography. Nat Commun 8:1349. doi: 10.1038/s41467-017-01285-x

Mandonnet E, Sarubbo S, Petit L (2018) The nomenclature of human white matter association pathways: proposal for a systematic taxonomic anatomical classification. Front Neuroanat 12:94. doi: 10.3389/fnana.2018.00094

Maunsell JH, van Essen DC (1983) The connections of the middle temporal visual area (MT) and their relationship to a cortical hierarchy in the macaque monkey. J Neurosci 3:2563–2586.

Meola A, Comert A, Yeh F-C, et al (2015) The controversial existence of the human superior fronto-occipital fasciculus: Connectome-based tractographic study with microdissection validation. Hum Brain Mapp 36:4964–4971. doi: 10.1002/hbm.22990

Meynert T (1892) Neue Studien über die Associations-Bündel des Hirnmantels. Math-Natural Sci Class 101:361–380.

Milchenko M, Marcus D (2013) Obscuring surface anatomy in volumetric imaging data. Neuroinformatics 11:65–75. doi: 10.1007/s12021-012-9160-3

Moeller S, Yacoub E, Olman CA, et al (2010) Multiband multislice GE-EPI at 7 tesla, with 16-fold acceleration using partial parallel imaging with application to high spatial and temporal whole-brain fMRI. Magn Reson Med 63:1144–1153. doi: 10.1002/mrm.22361

NITRC NITRC: MRIcron: Tool/resource Info. In: nitrc.org. https://www.nitrc.org/projects/mricron. Accessed 20 Apr 2020

Oishi H, Takemura H, Aoki SC, et al (2018) Microstructural properties of the vertical occipital fasciculus explain the variability in human stereoacuity. Proc Natl Acad Sci USA 115:12289–12294. doi: 10.1073/pnas.1804741115

Palejwala AH, O’Connor KP, Pelargos P, et al (2019) Anatomy and white matter connections of the lateral occipital cortex. Surg Radiol Anat. doi: 10.1007/s00276-019-02371-z

Panesar SS, Belo JTA, Yeh F-C, Fernandez-Miranda JC (2019) Structure, asymmetry, and connectivity of the human temporo-parietal aslant and vertical occipital fasciculi. Brain Struct Funct 224:907–923. doi: 10.1007/s00429-018-1812-0

Reveley C, Seth AK, Pierpaoli C, et al (2015) Superficial white matter fiber systems impede detection of long-range cortical connections in diffusion MR tractography. Proc Natl Acad Sci USA 112:E2820–8. doi: 10.1073/pnas.1418198112

Sachs H, Wernicke C (1892) Das Hemisphärenmark des menschlichen Grosshirns: Der Hinterhauptlappen. Das Hemisphärenmark des menschlichen Grosshirns: Der Hinterhauptlappen

Sampson JN, Wheeler WA, Yeager M, et al (2015) Analysis of Heritability and Shared Heritability Based on Genome-Wide Association Studies for Thirteen Cancer Types. J Natl Cancer Inst 107:djv279. doi: 10.1093/jnci/djv279

Schmahmann JD, Pandya DN (2007) The complex history of the fronto-occipital fasciculus. J Hist Neurosci 16:362–377. doi: 10.1080/09647040600620468

Schmahmann JD, Pandya DN (2006) Fiber pathways of the brain. doi: 10.1093/acprof:oso/9780195104233.001.0001

Schmahmann JD, Smith EE, Eichler FS, Filley CM (2008) Cerebral white matter: neuroanatomy, clinical neurology, and neurobehavioral correlates. Ann N Y Acad Sci 1142:266–309. doi: 10.1196/annals.1444.017

Schurr R, Filo S, Mezer AA (2019) Tractography delineation of the vertical occipital fasciculus using quantitative T1 mapping. Neuroimage 202:116121. doi: 10.1016/j.neuroimage.2019.116121

Setsompop K, Gagoski BA, Polimeni JR, et al (2012) Blipped-controlled aliasing in parallel imaging for simultaneous multislice echo planar imaging with reduced g-factor penalty. Magn Reson Med 67:1210–1224. doi: 10.1002/mrm.23097

Shibata S, Komaki Y, Seki F, et al (2015) Connectomics: comprehensive approaches for whole-brain mapping. Reprod Syst Sex Disord 64:57–67. doi: 10.1093/jmicro/dfu103

Shoja MM, Tubbs RS (2007) Augusta Déjerine-Klumpke: the first female neuroanatomist. Clin Anat 20:585–587. doi: 10.1002/ca.20474

Smith RE, Tournier J-D, Calamante F, Connelly A (2012) Anatomically-constrained tractography: improved diffusion MRI streamlines tractography through effective use of anatomical information. Neuroimage 62:1924–1938. doi: 10.1016/j.neuroimage.2012.06.005

Sotiropoulos SN, Moeller S, Jbabdi S, et al (2013) Effects of image reconstruction on fiber orientation mapping from multichannel diffusion MRI: reducing the noise floor using SENSE. Magn Reson Med 70:1682–1689. doi: 10.1002/mrm.24623

Sotiropoulos SN, Zalesky A (2019) Building connectomes using diffusion MRI: why, how and but. NMR Biomed 32:e3752. doi: 10.1002/nbm.3752

Takemura H, Pestilli F, Weiner KS (2018) Comparative neuroanatomy: Integrating classic and modern methods to understand association fibers connecting dorsal and ventral visual cortex. Neurosci Res. doi: 10.1016/j.neures.2018.10.011

Takemura H, Rokem A, Winawer J, et al (2016) A major human white matter pathway between dorsal and ventral visual cortex. Cereb Cortex 26:2205–2214. doi: 10.1093/cercor/bhv064

Thomas C, Ye FQ, Irfanoglu MO, et al (2014) Anatomical accuracy of brain connections derived from diffusion MRI tractography is inherently limited. Proc Natl Acad Sci USA 111:16574–16579. doi: 10.1073/pnas.1405672111

Tournier J-D, Calamante F, Connelly A (2012) MRtrix: Diffusion tractography in crossing fiber regions. Int J Imaging Syst Technol 22:53–66. doi: 10.1002/ima.22005

Tournier J-D, Calamante F, Connelly A (2007) Robust determination of the fibre orientation distribution in diffusion MRI: non-negativity constrained super-resolved spherical deconvolution. Neuroimage 35:1459–1472. doi: 10.1016/j.neuroimage.2007.02.016

Tournier J-D, Smith R, Raffelt D, et al (2019) MRtrix3: A fast, flexible and open software framework for medical image processing and visualisation. Neuroimage 202:116137. doi: 10.1016/j.neuroimage.2019.116137

Uesaki M, Takemura H, Ashida H (2018) Computational neuroanatomy of human stratum proprium of interparietal sulcus. Brain Struct Funct 223:489–507. doi: 10.1007/s00429-017-1492-1

Van Essen DC, Ugurbil K (2012) The future of the human connectome. Neuroimage 62:1299–1310. doi: 10.1016/j.neuroimage.2012.01.032

Van Essen DC, Ugurbil K, Auerbach E, et al (2012) The Human Connectome Project: a data acquisition perspective. Neuroimage 62:2222–2231. doi: 10.1016/j.neuroimage.2012.02.018

Vergani F, Mahmood S, Morris CM, et al (2014) Intralobar fibres of the occipital lobe: a post mortem dissection study. Cortex 56:145–156. doi: 10.1016/j.cortex.2014.03.002

Vialet N (1893) Les centres cérébraux de la vision et l’appareil nerveux visuel intra-cérébral. F. Alcan, Paris

Von Bonin GE, Garol HW, McCulloch WS (1942) The functional organization of the occipital lobe. Biol Symposia 7:165.

Wakana S, Jiang H, Nagae-Poetscher LM, et al (2004) Fiber tract-based atlas of human white matter anatomy. Radiology 230:77–87. doi: 10.1148/radiol.2301021640

Weiner KS, Yeatman JD, Wandell BA (2017) The posterior arcuate fasciculus and the vertical occipital fasciculus. Cortex 97:274–276. doi: 10.1016/j.cortex.2016.03.012

Wernicke C (1876) Das Urwindungssystem des menschlichen Gehirns. Archiv f Psychiatrie 6:298–326. doi: 10.1007/BF02230815

Wu Y, Sun D, Wang Y, et al (2016) Segmentation of the cingulum bundle in the human brain: A new perspective based on DSI tractography and fiber dissection study. Front Neuroanat 10:84. doi: 10.3389/fnana.2016.00084

Xu J, Strupp J, Auerbach EJ, et al (2012) Highly Accelerated Whole Brain Imaging Using Aligned-Blipped-Controlled-Aliasing Multiband EPI.

Yeatman JD, Weiner KS, Pestilli F, et al (2014) The vertical occipital fasciculus: a century of controversy resolved by in vivo measurements. Proc Natl Acad Sci USA 111:E5214–23. doi: 10.1073/pnas.1418503111

Zalesky A, Fornito A, Cocchi L, et al (2016) Connectome sensitivity or specificity: which is more important? Neuroimage 142:407–420. doi: 10.1016/j.neuroimage.2016.06.035

